# Tractography-based targeting of ventral intermediate nucleus: A comparison between conventional stereotactic targeting and diffusion tensor imaging-based targeting

**DOI:** 10.1101/2021.04.29.441001

**Authors:** Anupa A. Vijayakumari, Drew Parker, Andrew I Yang, Ashwin G. Ramayya, Ronald L. Wolf, Darien Aunapu, Gordon H. Baltuch, Ragini Verma

**Affiliations:** DiCIPHR (Diffusion and Connectomics in Precision Healthcare Research) Lab, Department of Radiology, Perelman School of Medicine, University of Pennsylvania, Philadelphia, Pennsylvania, USA; Department of Neurosurgery, Perelman School of Medicine, University of Pennsylvania, Philadelphia, Pennsylvania, USA; Department of Radiology, Perelman School of Medicine, University of Pennsylvania, Philadelphia, Pennsylvania, USA

**Keywords:** deep brain stimulation, dentato-rubro-thalamic tract, deterministic tractography, diffusion tensor imaging tractography, magnetic resonance imaging guided focused ultrasound, probabilistic tractography, ventral intermediate nucleus

## Abstract

**Background:** The ventral intermediate (VIM) nucleus of the thalamus is the main target for lesioning using magnetic resonance imaging (MRI) guided focused ultrasound (MRgFUS) or deep brain stimulation (DBS). Targeting of VIM still depends on standard stereotactic coordinates, which do not account for inter-individual variability. Several approaches have been proposed including visualization of dentato-rubro-thalamic tract (DRTT) using diffusion tensor imaging tractography.

**Objective:** To compare probabilistic tracking of DRTT with deterministic tracking of DRTT and stereotactic coordinates to identify the most appropriate approach to target VIM.

**Methods:** In this retrospective study, we assessed the VIM targeted using stereotactic coordinates, deterministic and probabilistic tracking of DRTT in 19 patients with essential tremor who underwent DBS with VIM targeted using microelectrode recordings. We subsequently determined the positions of VIM derived from these three approaches and compared with that of DBS lead position using paired sample *t*-tests.

**Results:** The probabilistic tracking of DRTT was significantly anterior to the lead (1.45 ± 1.61 mm (*P*< 0.0001)), but not in the medial/lateral position (−0.29±2.42 mm (*P*=0.50)). Deterministic tracking of DRTT was significantly lateral (2.16 ± 1.94 mm (*P*< 0.0001)) and anterior to the lead (1.66 ± 2.1 mm (*P*< 0.0001)). The stereotactic coordinates were significantly lateral (2.41 ± 1.41 mm (*P*< 0.0001)) and anterior (1.23 ± 0.89 mm (*P*< 0.0001)) to the lead.

**Conclusion:** Probabilistic tracking of DRTT was found to be superior in targeting VIM compared to deterministic tracking and stereotactic coordinates.

## 1. Introduction

The goal of this paper is to provide a tractography-based localization of ventral intermediate (VIM) nucleus of the thalamus, which is crucial for magnetic resonance imaging (MRI) guided focused ultrasound (MRgFUS) or deep brain stimulation (DBS) in the treatment of medically refractory essential tremor (ET) and tremor-dominant Parkinson’s disease.^1^ VIM is primarily involved in sending and receiving electric impulses to and from the motor cortex and the cerebellum to control movement^2^, and hence targeted for lesioning using MRgFUS or DBS to alleviate tremor. VIM is a relatively small anatomic structure (4mm×4mm×6mm) which cannot be readily identified on structural MRI. Therefore, VIM targeting is often performed indirectly using stereotactic coordinates referencing the mid-commissural line and third ventricle. However, there are no universally agreed upon stereotactic coordinates, with different targeting formulas used across centers. ^3^ Generally, VIM is targeted approximately 14-15 mm lateral to the midline (or 11 mm lateral from the wall of the third ventricle), and 25% of length of the anterior commissure (AC)–posterior commissure (PC) line anterior to the PC, at the level of the commissural plane.^3-6^ Although this method provides an approximation of VIM location, final implant positions are determined with intraoperative microelectrode recordings (MER) and neurologic testing with stimulation to verify the target region and identify its borders. However, MER is time consuming and cannot be readily applied in the setting of asleep DBS and incisionless procedures such as MRgFUS^7,8^.

In recent years, visualization of the dentato-rubro-thalamic tract (DRTT) using diffusion tensor imaging (DTI) tractography has been used for identifying VIM for DBS and MRgFUS.^9-11^ According to the classical definition, DRTT has a decussating pathway (d-DRTT), which emanates from the dentate nucleus (DN), passing through the superior cerebellar peduncle (SCP), crossing the midline to the contralateral red nucleus (RN) and VIM, where it synapses with neurons ascending to the motor cortex. These anatomical courses were defined first in monkeys^12,13^ and later in humans using post-mortem and DTI tractography studies.^14-16^ In addition to the d-DRTT, a non-decussating pathway of DRTT (nd-DRTT) connecting to the ipsilateral RN and the thalamus has also been reported using microdissection and DTI studies in humans.^17^ Although the anatomy of DRTT has been reported through RN, the course of DRTT through the subthalamic area has also been reported in humans.^18-20^

A variety of ways of tracking the DRTT have been reported. Several approaches use regions of interest (ROIs), including DN, RN or posterior subthalamic area (PSA), and the motor cortex or precentral gyrus. ^10,20-25^ Alternately, Sammartino et al tracked the pyramidal tract (PT) and medial lemniscus (MLm) to locate a seed region to track the DRTT.^26^ Most studies examining DRTT for VIM planning applied deterministic fiber tracking algorithms because of its ease of analysis and seamless integration with the stereotactic targeting software for DBS surgery or MRgFUS.^5,8,27-29^ However, deterministic DRTT (detDRTT) is inaccurate in crossing fiber regions or in obtaining decussating pathways^5,26^. Probabilistic tracking using higher-order diffusion models can overcome crossing fiber issues, and has been reported to be superior in identifying DRTT.^30^ However, there is a dearth of research examining the VIM positions derived from probDRTT.

Accurate initial targeting of VIM is important for favorable patient outcome and efficacy of the DBS and MRgFUS procedures. VIM targeting using stereotactic coordinates is prone to anatomical variability in third ventricular widths and AC-PC lengths, leading to targeting error^31^. Although VIM targeting based on tractography of DRTT can accommodate these anatomical variations, precision of detDRTT does not always precisely coincide with DBS or MRgFUS targeting locations^7,9^ and also fails to consistently create d-DRTT and/or nd-DRTT^23,26^. In this study, we propose a new protocol for VIM targeting utilizing probabilistic tractography, which can consistently reconstruct d-DRTT and nd-DRTT. This is compared to VIM targeting based on stereotactic coordinates, as well as deterministic tractography using the Sammartino method.^26^ This is evaluated in a retrospective study in patients with ET, who underwent standard awake DBS using intraoperative MER, which is the gold-standard for VIM localization to implant DBS leads.

## 2. Methods

### 2.1. Patient cohort

We studied 19 patients who underwent DBS surgery for ET (M:F=10:9), mean age (±SD)=66.05 (±12.07) years. 13 patients had bilateral implants and 6 patients had unilateral implants (2 right hemisphere and 4 left hemisphere), allowing investigation of 32 DBS leads. Patient selection, DBS surgery procedure, and ethical approval have already been described in detail.^5^ Briefly, patients underwent stereotactic placement of DBS leads with a Leksell frame (Elekta, Stockholm, Sweden) and VIM was targeted indirectly in the commissural plane as follows: medial-lateral (ML) coordinates; 14 mm lateral to the midline or 11 mm from the lateral wall of third ventricle, anterior-posterior (AP) coordinates; 25% of AC-PC distance from the PC. Yang et al, previously evaluated proximity of the detDRTT to DBS stimulation parameters in a cohort of 26 patients with ET^5^. For this study, 7 patients were excluded due to unavailability of whole brain T1-weighted image (n=1), incomplete DTI/FreeSurfer processing (n=4) and use of different DTI scanning parameters (n=2), leading to 19 patients with ET.

### 2.2. MRI acquisition

Magnetic Resonance Imaging (MRI) scans were acquired on a 3T GE Discovery MR750w scanner using a 19-channel head coil (GE Healthcare, Waukesha Wisconsin, USA). The following sequences were acquired: 1) T1 SPGR structural MRI scan with TR/TE=6.3/1.46 ms, flip angle=20°, slice thickness=2 mm, reconstructed every 1 mm, 320×256 acquisition matrix, and in-plane resolution of 0.43×0.43 mm, and 2) DTI data with 1 b=0 s/mm^2^, 33 gradient directions with b-value=2000 s/mm^2^, TR/TE=14s/0.113 s, field of view 26 cm, 128×128 acquisition matrix, slice thickness=3 mm, and in-plane resolution of 1×1 mm.

### 2.3. Fiber Tracking Technique

#### 2.3.1. Deterministic fiber tracking of DRTT

DetDRTT was performed using the method of Sammartino et al^26^, which uses PT and MLm as internal landmarks. Anatomically, PT and MLm represent, respectively, the lateral and the posterior ventral boundaries of the VIM of thalamus. A thalamic ROI with its center equidistant (3 mm) from the borders of the PT and MLm was placed at the AC-PC level to reconstruct the DRTT. DynaSuite Neuro (InVivo Corp., Pewaukee, Wisconsin, USA) was used for tractography with previously described parameters.^5^

#### 2.3.2. Probabilistic Fiber Tractography of DRTT

DTI datasets were denoised^32^ and corrected for eddy currents and motion with *eddy*^33^ in FSL^34^. The T1-weighted structural images were bias-corrected using N4BiasCorrection tool from Advanced Normalization Tools (ANTs)^35^ and skull stripping was achieved using multi-atlas region segmentation utilizing ensembles of registration algorithms and parameters (MUSE)^36^. DTI were resliced to 2×2×2 mm and T1 images were coregistered to the DTI data with ANTs.^35^ Constrained spherical deconvolution was performed to estimate the fiber orientation distribution (FOD) in each voxel using the MRtrix3 package.^37^ Then probabilistic tractography was performed using the second-order Integration over Fiber Orientation Distributions (iFOD2) algorithm.^38^ The seed region for the reconstruction of DRTT was the thalamus and the target ROIs were hand motor region and bilateral DN. DN and hand motor region were hand-drawn on the b0 and fractional anisotropy (FA) images respectively; left and right thalamus were automatically segmented using FreeSurfer^39^ on T1 weighted images, as shown in Fig. 1(a). Exclusion ROIs included internal capsule at the level of AC-PC, ventral part of the brainstem to prevent the tracking algorithm from merging with the PT or MLm, and corpus callosum (Figure 1(b)). 500 seeds were placed at random in each voxel of the thalamus, keeping streamlines connecting target ROIs. Tracking parameters were a minimum fiber length=20 mm, maximum fiber length=250 mm, step size=0.5 mm, and angle threshold=45°. Figure 2 shows the probDRTT in one patient.

**Figure 1:**
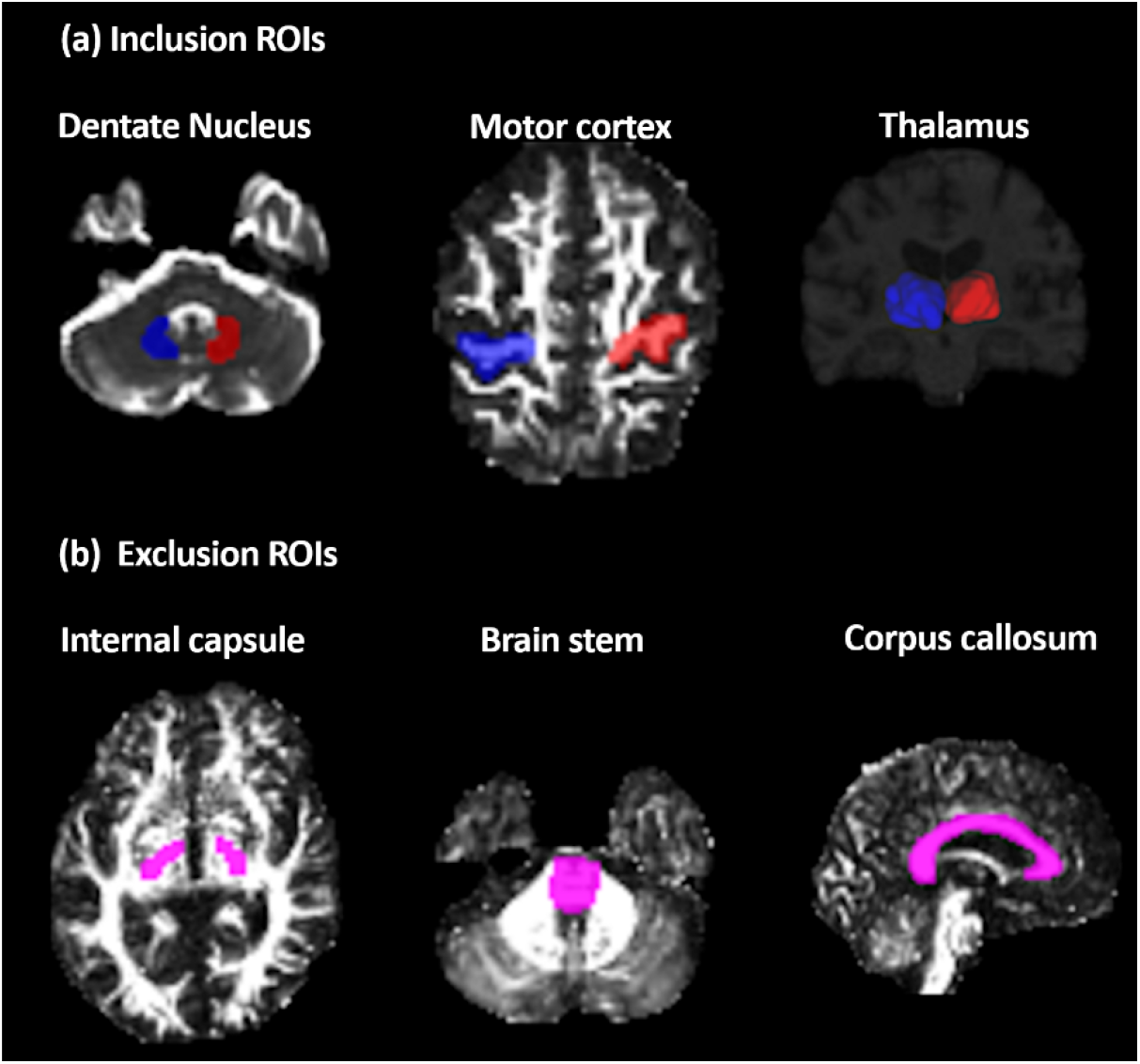
Delineation of the (a) inclusion regions of interests (dentate nucleus, motor cortex and thalamus) and (b) exclusion ROIs (internal capsule at the level of AC-PC) for the reconstruction of dentato-rubro-thalamic tract. Blue color indicates the ROIs on the right side of the brain, red color indicates the ROIs on the left side of the brain, and pink color indicates the exclusion masks.

**Figure 2.**
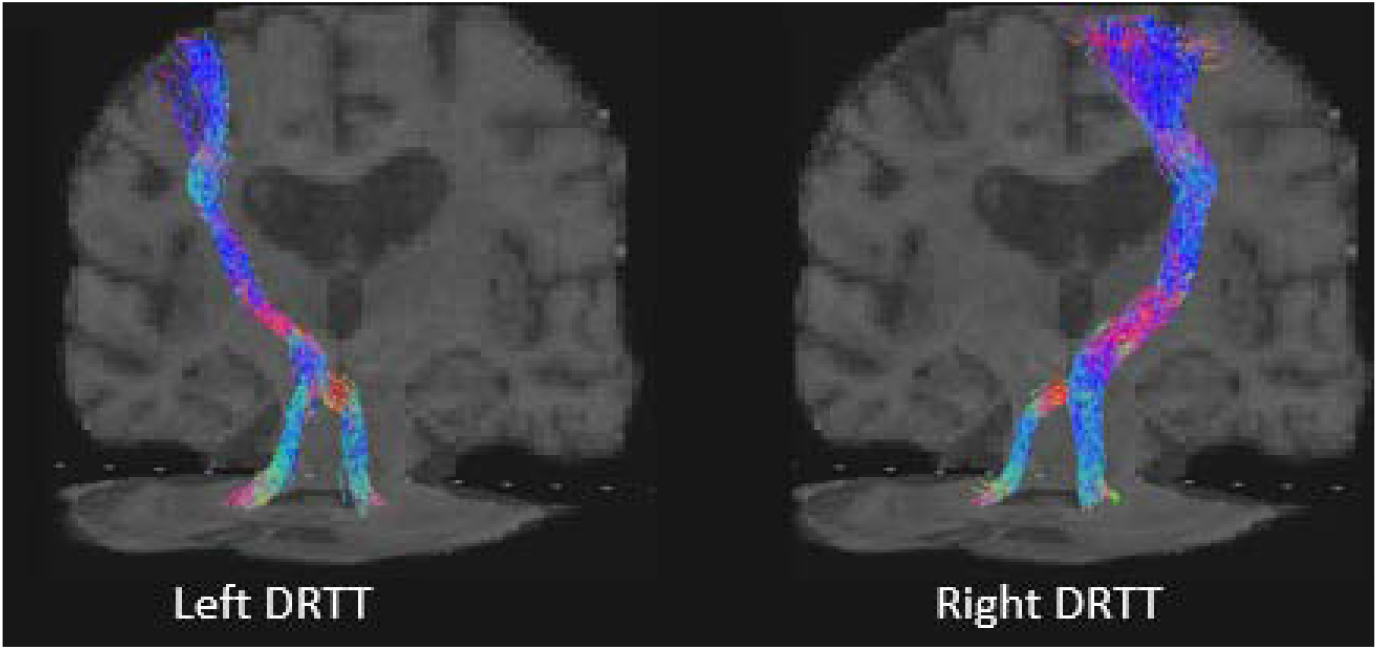
Representation of dentato-rubro-thalamic tract in one patient with essential tremor

### 2.4. Calculation of centroids

The T1 images of the patients were manually AC-PC aligned in ITK-Snap software^40^, and rigidly registered to the FA maps using ANTs. The track-density image (TDI) of the DRTT was obtained^41^ and transformed to AC-PC aligned T1 space and inspected within the axial slice containing the AC-PC line. The center of gravity (centroid) within this plane was calculated by using the TDI values as weights and transformed to medial/lateral (ML) and anterior/posterior (AP) coordinates by subtracting the real-world coordinates of the PC.

### 2.5. Clinical assessments

Tremor was assessed in a subset of patients (n=10) before and after the DBS surgery using the Clinical Rating Scale for Tremor (CRST) score (ranging from 0–160 points, higher scores indicating greater disability).^42^

### 2.6. Comparison between different methods of VIM localization

Paired sample *t*-tests were performed between AP and ML positions of lead center to the centers of stereotactic coordinates, Sammartino-based detDRTT and probDRTT. A *P* value of 0.05 was considered statistically significant. All analysis was performed in SPSS version 21.

## 3. Results

The demographics and clinical characteristics of the patients are shown in Table 1.

**Table 1.**
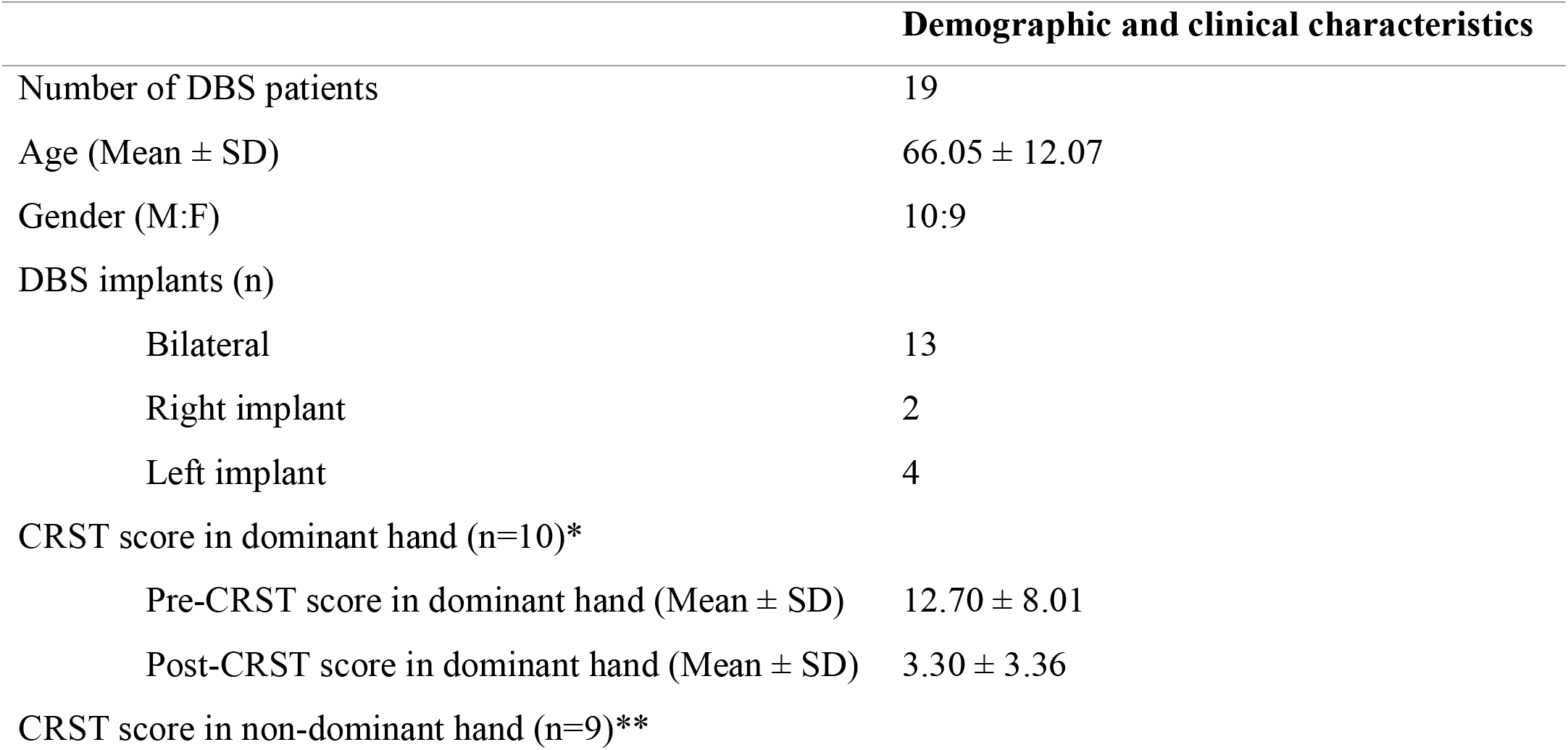
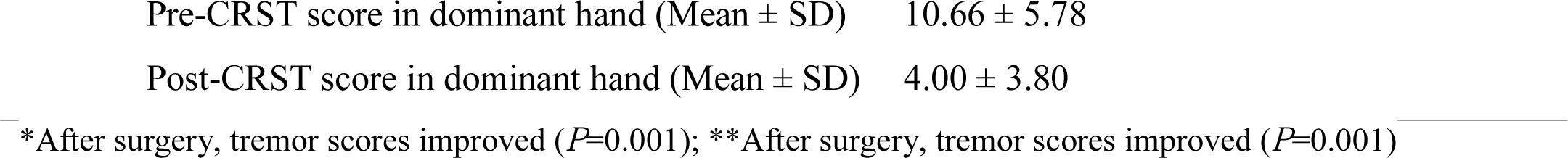
Demographic and clinical characteristics of the patients.

Of 32 hemispheres studied, nd-DRTT and d-DRTT traversing to ipsilateral motor cortical output fibers were demonstrated in all hemispheres using probDRT. Using Sammartino-based detDRTT, although nd-DRTT was reconstructed in all hemispheres, d-DRTT was reconstructed only in 11 hemispheres (31%).

At the level of the intercommissural line, leads were located 12.05 ± 1.35 mm lateral to the midline and 5.19 ± 0.91 mm anterior to the PC. The probDRTT was located 11.76 ± 2.04 mm lateral to the midline and 6.64 ± 1.64 mm anterior to the PC. The detDRTT was located 14.22 ± 2.27 mm lateral to the midline and 6.85 ± 2.21 mm anterior to the PC. The stereotactic coordinates were located 14.46 ± 0.59 mm lateral to the midline and 6.42 ± 0.44 mm anterior to the PC (Table 2).

**Table 2.** Deep Brain Stimulation Lead and Dentato-rubro-thalamic Tract Coordinates

|  | Lateral distance from AC-PC<br>(mm) | Anterior distance from AC-PC<br>(mm) |
| --- | --- | --- |
| DBS leads | 12.05 ± 1.35 | 5.19 ± 0.91 |
| ProbDRTT | 11.76 ± 2.04 | 6.64 ± 1.64 |
| Sammartino-based detDRTT | 14.22 ± 2.27 | 6.85 ± 2.21 |
| Stereotactic coordinates | 14.46 ± 0.59 | 6.42 ± 0.44 |

Hence, the probDRTT was significantly anterior to the lead by 1.45 ± 1.61 mm (*P*<0.0001), but not significant in the ML position -0.29 ± 2.42 mm (*P*= 0.50). The detDRTT was significantly lateral and anterior to the lead, with a distance of 2.16 ± 1.94 mm (*P*<0.0001) and 1.66 ± 2.1 mm (*P*<0.0001) respectively. The stereotactic coordinates were also significantly lateral and anterior to the lead, with a distance of 2.41 ± 1.41 mm (*P*<0.0001) and 1.23 ± 0.89 mm (*P*<0.0001) respectively. Figure 3 depicts the coordinates of the centers of DBS lead, probDRTT, Sammartino-based detDRTT, and stereotactic coordinates in the commissural plane.

**Figure 3.**
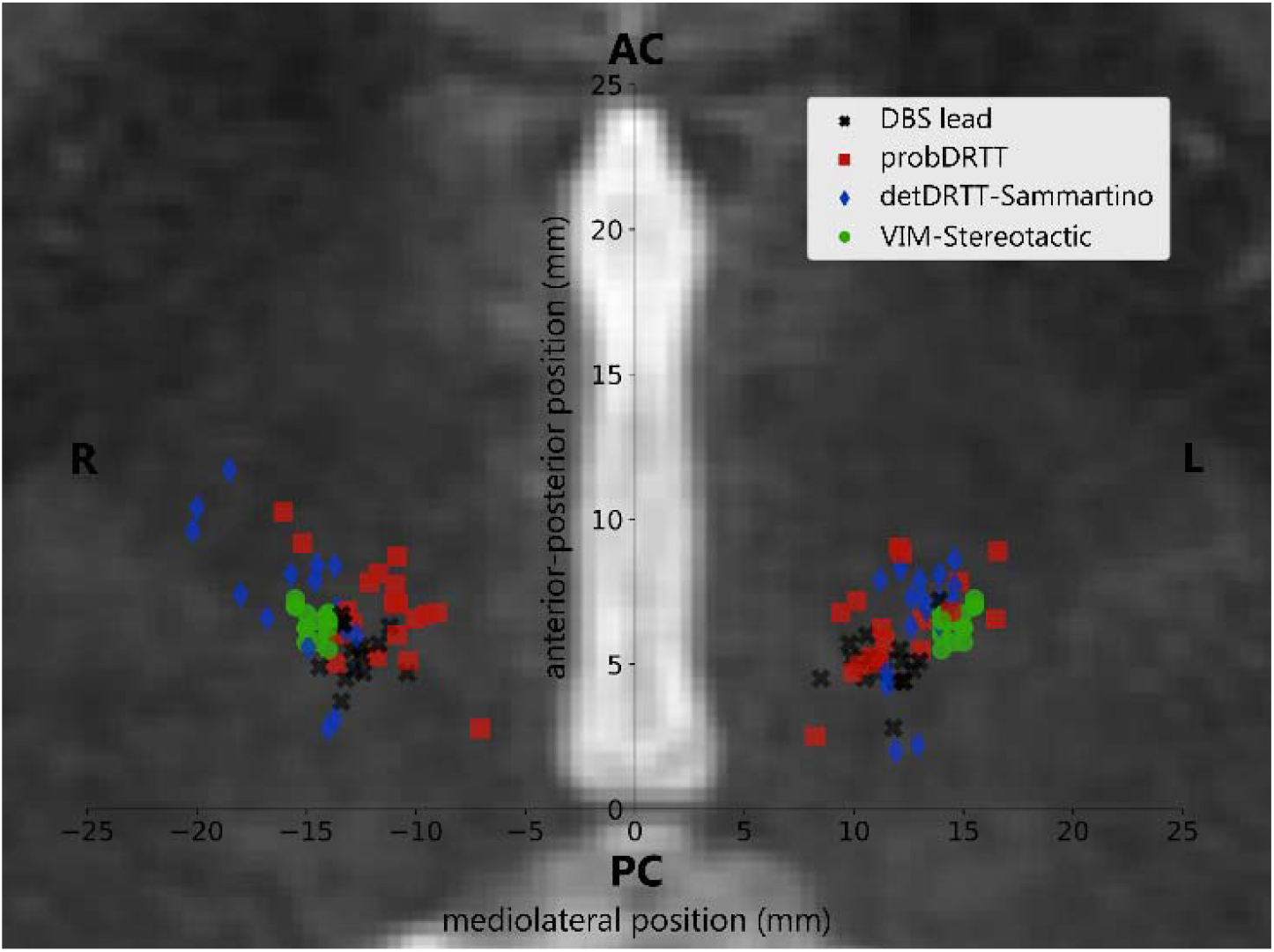
Coordinates of the centers of DBS lead, probDRTT, Sammartino-based detDRTT, and stereotactic coordinates in the commissural plane.

## 4. Discussion

Precise targeting of the VIM is important for favorable patient outcome and efficacy of DBS and MRgFUS procedures. In this study, we demonstrated better VIM localization using probabilistic tracking compared to deterministic tracking, and stereotactic coordinates based on their comparison with DBS lead positions using MER. VIM targeted using probDRTT was anterior to the DBS leads with no medial or lateral bias, while stereotactic coordinates and detDRTT were significantly anterior and lateral to the DBS leads.

Tractography has been widely used for VIM targeting by neurosurgeons for preoperative planning.^43^ Currently, the only method available in commercial neurosurgical planning software is deterministic tractography. While DRTT is a complex cerebello-thalamic tract with decussating and non-decussating pathways,^14,17^ deterministic tractography is insensitive in detecting crossing, kissing or other complex fiber arrangements,^44,45^with a low success rate in reconstructing DRTT consistently across patients.^5,23,26^ Few studies have compared deterministic and probabilistic tractography, with detDRTT being less sensitive in DRTT identification. Schlaier et al., reported that deterministic tractography was less sensitive compared to probabilistic tractography for identifying d-DRTT^30^. Sammartino et al. successfully reconstructed d-DRTT and nd-DRTT using probabilistic tractography. Although deterministic tractography produced nd-DRTT, d-DRTT was identified only in 62% of cases studied.^26^ When using the Sammartino method for detDRTT we observed that nd-DRTT was consistently reconstructed but not d-DRTT. On the other hand, probabilistic tractography successfully generated non-decussating and decussating DRTT. Thus, we demonstrate that probabilistic tractography is successful in identifying the decussating and non-decussating pathways of the DRTT unlike deterministic tracking.

Further, we studied the VIM locations generated using stereotactic coordinates, detDRTT and probDRTT in the commissural plane. Yang et al.^5^ who used Sammartino-based detDRTT in 26 patients with DBS observed that DRTT was anterior to the lead by 1.5 and lateral by 2.1 mm, which is consistent with present study in a subset of 19 patients with DBS, where Sammartino-based detDRTT was 1.6 mm anterior and 2.1 lateral to the lead. In the original study by Sammartino et al.,^26^ DRTT was 1.8 mm anterior and 1 mm lateral to the leads in 18 patients with ET. Another study by the same group using probabilistic tractography in a cohort of patients with focused ultrasound, found VIM location determined by probDRTT to be 0.9 mm anterior to the lesion^29^. In our study, probDRTT was significantly anterior to the lead by 1.4 mm with no medio-lateral bias. Finally, the VIM stereotactic coordinates were significantly anterior and lateral to the lead. Our results suggest that centroids of probDRTT as the initial target are superior to using the stereotactic coordinates or detDRTT in targeting VIM. However, the results should be interpreted cautiously because of the anterior displacement of probDRTT. It is unclear whether this is rooted in individual variability or a reflection of true difference. In future, studies in a larger sample size would be necessary to tease this apart.

There are some limitations to our study. Sample size of this study is small, and in future the probDRTT results will need to be validated in a larger cohort of patients for VIM targeting. Until now, only detDRTT have been integrated into surgical planning platforms, and probDRTT is not approved for clinical practice. Despite these limitations, we believe that our findings support further investigation of the use of probDRTT to consistently produce the DRTT for the localization of VIM for DBS and MRgFUS.

## 5. Conclusions

We observed that VIM targeted using stereotactic coordinates and Sammartino-based detDRTT were anterior and lateral to the lead. ProbDRTT was anterior to the lead but did not have any bias in the medial-lateral direction. Our results suggest that probDRTT is superior to stereotactic coordinates and detDRTT in targeting the VIM for DBS or MRgFUS procedures.

## References

1. Benabid AL, Pollak P, Gervason C, et al. Long-term suppression of tremor by chronic stimulation of the ventral intermediate thalamic nucleus. Lancet. 1991;337(8738):403–406.

2. Dormont D, Cornu P, Pidoux B, et al. Chronic thalamic stimulation with three-dimensional MR stereotactic guidance. AJNR Am J Neuroradiol. 1997;18(6):1093–1107.

3. Papavassiliou E, Rau G, Heath S, et al. Thalamic deep brain stimulation for essential tremor: relation of lead location to outcome. Neurosurgery. 2004;54(5):1120-1129; discussion 1129-1130.

4. King NKK, Krishna V, Sammartino F, et al. Anatomic Targeting of the Optimal Location for Thalamic Deep Brain Stimulation in Patients with Essential Tremor. World Neurosurg. 2017;107:168–174.

5. Yang AI, Buch VP, Heman-Ackah SM, et al. Thalamic deep brain stimulation for essential tremor: relation of the dentato-rubro-thalamic tract with stimulation parameters. World Neurosurg. 2020.

6. Osenbach RK, Burchiel R. Thalamotomy: indications, techniques, and results. Park Ridge, IL: American Association of Neurological Surgeons; 1998: 107–129.

7. Miller TR, Zhuo J, Eisenberg HM, et al. Targeting of the dentato-rubro-thalamic tract for MR-guided focused ultrasound treatment of essential tremor. The neuroradiology journal. 2019;32(6):401–407.

8. Coenen VA, Sajonz B, Prokop T, et al. The dentato-rubro-thalamic tract as the potential common deep brain stimulation target for tremor of various origin: an observational case series. Acta Neurochir (Wien). 2020;162(5):1053–1066.

9. Anthofer J, Steib K, Fellner C, Lange M, Brawanski A, Schlaier J. The variability of atlas-based targets in relation to surrounding major fibre tracts in thalamic deep brain stimulation. Acta Neurochir (Wien). 2014;156(8):1497-1504; discussion 1504.

10. Schlaier J, Anthofer J, Steib K, et al. Deep brain stimulation for essential tremor: targeting the dentato-rubro-thalamic tract? Neuromodulation : journal of the International Neuromodulation Society. 2015;18(2):105–112.

11. Chazen JL, Sarva H, Stieg PE, et al. Clinical improvement associated with targeted interruption of the cerebellothalamic tract following MR-guided focused ultrasound for essential tremor. Journal of neurosurgery. 2018;129(2):315–323.

12. Rouiller EM, Liang F, Babalian A, Moret V, Wiesendanger M. Cerebellothalamocortical and pallidothalamocortical projections to the primary and supplementary motor cortical areas: a multiple tracing study in macaque monkeys. J Comp Neurol. 1994;345(2):185–213.

13. Asanuma C, Thach WR, Jones EG. Anatomical evidence for segregated focal groupings of efferent cells and their terminal ramifications in the cerebellothalamic pathway of the monkey. Brain Res. 1983;286(3):267–297.

14. Kwon HG, Hong JH, Hong CP, Lee DH, Ahn SH, Jang SH. Dentatorubrothalamic tract in human brain: diffusion tensor tractography study. Neuroradiology. 2011;53(10):787–791.

15. Palesi F, Tournier JD, Calamante F, et al. Contralateral cerebello-thalamo-cortical pathways with prominent involvement of associative areas in humans in vivo. Brain Struct Funct. 2015;220(6):3369–3384.

16. Mollink J, van Baarsen KM, Dederen PJ, et al. Dentatorubrothalamic tract localization with postmortem MR diffusion tractography compared to histological 3D reconstruction. Brain Struct Funct. 2016;221(7):3487–3501.

17. Meola A, Comert A, Yeh FC, Sivakanthan S, Fernandez-Miranda JC. The nondecussating pathway of the dentatorubrothalamic tract in humans: human connectome-based tractographic study and microdissection validation. Journal of neurosurgery. 2016;124(5):1406–1412.

18. Gallay MN, Jeanmonod D, Liu J, Morel A. Human pallidothalamic and cerebellothalamic tracts: anatomical basis for functional stereotactic neurosurgery. Brain Struct Funct. 2008;212(6):443–463.

19. Morel A, Magnin M, Jeanmonod D. Multiarchitectonic and stereotactic atlas of the human thalamus. J Comp Neurol. 1997;387(4):588–630.

20. Nowacki A, Schlaier J, Debove I, Pollo C. Validation of diffusion tensor imaging tractography to visualize the dentatorubrothalamic tract for surgical planning. Journal of neurosurgery. 2018;130(1):99–108.

21. Akram H, Dayal V, Mahlknecht P, et al. Connectivity derived thalamic segmentation in deep brain stimulation for tremor. Neuroimage Clin. 2018;18:130–142.

22. Hana A, Hana A, Dooms G, Boecher-Schwarz H, Hertel F. Depiction of dentatorubrothalamic tract fibers in patients with Parkinson’s disease and multiple sclerosis in deep brain stimulation. BMC research notes. 2016;9:345.

23. Nowacki A, Debove I, Rossi F, et al. Targeting the posterior subthalamic area for essential tremor: proposal for MRI-based anatomical landmarks. Journal of neurosurgery. 2018;131(3):820–827.

24. Coenen VA, Allert N, Madler B. A role of diffusion tensor imaging fiber tracking in deep brain stimulation surgery: DBS of the dentato-rubro-thalamic tract (drt) for the treatment of therapy-refractory tremor. Acta Neurochir (Wien). 2011;153(8):1579-1585; discussion 1585.

25. Coenen VA, Allert N, Paus S, Kronenburger M, Urbach H, Madler B. Modulation of the cerebello-thalamo-cortical network in thalamic deep brain stimulation for tremor: a diffusion tensor imaging study. Neurosurgery. 2014;75(6):657-669; discussion 669-670.

26. Sammartino F, Krishna V, King NK, et al. Tractography-Based Ventral Intermediate Nucleus Targeting: Novel Methodology and Intraoperative Validation. Mov Disord. 2016;31(8):1217–1225.

27. Dembek TA, Petry-Schmelzer JN, Reker P, et al. PSA and VIM DBS efficiency in essential tremor depends on distance to the dentatorubrothalamic tract. Neuroimage Clin. 2020;26:102235.

28. Coenen VA, Varkuti B, Parpaley Y, et al. Postoperative neuroimaging analysis of DRT deep brain stimulation revision surgery for complicated essential tremor. Acta Neurochir (Wien). 2017;159(5):779–787.

29. Krishna V, Sammartino F, Agrawal P, et al. Prospective Tractography-Based Targeting for Improved Safety of Focused Ultrasound Thalamotomy. Neurosurgery. 2019;84(1):160–168.

30. Schlaier JR, Beer AL, Faltermeier R, et al. Probabilistic vs. deterministic fiber tracking and the influence of different seed regions to delineate cerebellar-thalamic fibers in deep brain stimulation. The European journal of neuroscience. 2017;45(12):1623–1633.

31. Brierley JB, Beck E. The significance in human stereotactic brain surgery of individual variation in the diencephalon and globus pallidus. J Neurol Neurosurg Psychiatry. 1959;22:287–298.

32. Manjon JV, Coupe P, Concha L, Buades A, Collins DL, Robles M. Diffusion weighted image denoising using overcomplete local PCA. PloS one. 2013;8(9):e73021.

33. Andersson JLR, Sotiropoulos SN. An integrated approach to correction for off-resonance effects and subject movement in diffusion MR imaging. Neuroimage. 2016;125:1063–1078.

34. Smith SM, Jenkinson M, Woolrich MW, et al. Advances in functional and structural MR image analysis and implementation as FSL. Neuroimage. 2004;23(S1):208–219.

35. Avants BB, Tustison NJ, Johnson HJ. ANTs Distribution, http://stnava.github.io/ANTs/. http://stnava.github.io/ANTs/.

36. Doshi J, Erus G, Ou Y, et al. MUSE: MUlti-atlas region Segmentation utilizing Ensembles of registration algorithms and parameters, and locally optimal atlas selection. NeuroImage. 2016;127:186–195.

37. Tournier J-D, Calamante F, Connelly A. MRtrix: Diffusion tractography in crossing fiber regions. International Journal of Imaging Systems and Technology. 2012;22:53–66.

38. Tournier J-D CF, Connelly A. Improved probabilistic streamlines tractography by 2nd order integration over fibre orientation distributions. Paper presented at: Proceedings of the International Society for Magnetic Resonance in Medicine. 2010.

39. Fischl B, van der Kouwe A, Destrieux C, et al. Automatically parcellating the human cerebral cortex. Cerebral cortex. 2004;14(1):11–22.

40. Yushkevich PA, Piven J, Hazlett HC, et al. User-guided 3D active contour segmentation of anatomical structures: significantly improved efficiency and reliability. Neuroimage. 2006;31(3):1116–1128.

41. Dhollander T, Emsell L, Van Hecke W, Maes F, Sunaert S, Suetens P. Track orientation density imaging (TODI) and track orientation distribution (TOD) based tractography. Neuroimage. 2014;94:312–336.

42. Stacy MA, Elble RJ, Ondo WG, Wu SC, Hulihan J, group TRSs. Assessment of interrater and intrarater reliability of the Fahn-Tolosa-Marin Tremor Rating Scale in essential tremor. Mov Disord. 2007;22(6):833–838.

43. Gravbrot N, Saranathan M, Pouratian N, Kasoff WS. Advanced Imaging and Direct Targeting of the Motor Thalamus and Dentato-Rubro-Thalamic Tract for Tremor: A Systematic Review. Stereotact Funct Neurosurg. 2020;98(4):220–240.

44. Calamante F. The Seven Deadly Sins of Measuring Brain Structural Connectivity Using Diffusion MRI Streamlines Fibre-Tracking. Diagnostics (Basel). 2019;9(3).

45. Jbabdi S, Johansen-Berg H. Tractography: where do we go from here? Brain Connect. 2011;1(3):169–183.

